# A Zika virus protein expression screen in *Drosophila* to investigate targeted host pathways during development

**DOI:** 10.1101/2023.04.28.538736

**Authors:** Nichole Link, J Michael Harnish, Brooke Hull, Shelley Gibson, Miranda Dietze, Uchechukwu E. Mgbike, Silvia Medina-Balcazar, Priya S. Shah, Shinya Yamamoto

## Abstract

In the past decade, Zika virus (ZIKV) emerged as a global public health concern. While adult infections are typically mild, maternal infection can lead to adverse fetal outcomes. Understanding how ZIKV proteins disrupt development can provide insights into the molecular mechanisms of symptoms caused by this virus including microcephaly. In this study, we generated a toolkit to ectopically express Zika viral proteins *in vivo* in *Drosophila melanogaster* in a tissue-specific manner using the GAL4/UAS system. We use this toolkit to identify phenotypes and host pathways targeted by the virus. Our work identified that expression of most ZIKV proteins cause scorable phenotypes, such as overall lethality, gross morphological defects, reduced brain size, and neuronal function defects. We further use this system to identify strain-dependent phenotypes that may contribute to the increased pathogenesis associated with the more recent outbreak of ZIKV in the Americas. Our work demonstrates *Drosophila’s* use as an efficient *in vivo* model to rapidly decipher how pathogens cause disease and lays the groundwork for further molecular study of ZIKV pathogenesis in flies.

## INTRODUCTION

Zika virus (ZIKV) is a small RNA virus that gained notoriety for its ability to cause a spectrum of birth defects dubbed Congenital Zika Syndrome (CZS) (da Silva Pone et al., 2018). This disorder occurs in infants after infection *in utero* and can include microcephaly, ocular abnormalities, congenital contractures, and hypertonia restricting body movements, with the possibility of additional defects arising later in development (Medina and Medina-Montoya, 2017). Some children without physical findings at birth also develop adverse neurological defects, developmental delays, and microcephaly at a later point (Aragao et al., 2017; Bertolli et al., 2020; Carvalho et al., 2019; Rice et al., 2018; van der Linden et al., 2016), indicating that ZIKV has much broader effect on neurodevelopment and function than initially appreciated. Microcephaly, or reduced head size typically associated with smaller brain size, is one of the most severe outcomes in CZS and is associated with cognitive and neurological defects. Given the severe outcomes associated with ZIKV infection *in utero*, it is critical to understand how ZIKV disrupts brain development.

In adults, ZIKV infection can be mild with rash, fever, or muscle pain. However, some patients develop Guillain-Barré syndrome, an autoimmune defect where the body attacks its own peripheral nervous system. Patients affected by Guillain-Barré syndrome often experience muscle weakness or severe paralysis, but most people recover from this condition (Cao-Lormeau et al., 2016). In addition, ZIKV has also been linked to severe neurological disease in adults (Mehta et al., 2018). This includes meningoencephalitis (Carteaux et al., 2016), sensory neuropathy (Medina et al., 2016), and seizures (Asadi-Pooya, 2016). Since the spectrum of disease is wide, a greater understanding of how ZIKV affects mature tissues in addition to developing tissues would be beneficial for patient treatment.

In addition to clinical evidence, animal models have also shown that ZIKV can have long term impact on nervous system function. For example, ZIKV infections in mice led to long term neuropathological effects that persisted into adulthood (Nem de Oliveira Souza et al., 2018), and postnatal infections in rhesus macaques resulted in behavioral, motor, and cognitive deficits associated with abnormalities in brain structure (Raper et al., 2020). Thus, it is likely this pathogen affects multiple molecular pathways that are associated with development and neural function, resulting in the wide spectrum of phenotypes associated with ZIKV infection during development or in adulthood.

The mechanism by which ZIKV causes disease, especially microcephaly, has become a major research topic since the declaration of ZIKV epidemic in 2016. Current data suggests that molecular mechanisms of ZIKV-induced microcephaly may be mediated by inhibition of multiple host pathways by ZIKV proteins. The genome of ZIKV is ∼10kb and encodes three structural proteins [Capsid (C), precursor Membrane (prM) and Envelope (E)] and seven nonstructural proteins (NS1, NS2A, NS2B, NS3, NS4A, NS4B and NS5) (**Figure 1A**). In one study, Liang *et al*. found that ZIKV NS4A and NS4B suppress Akt-mTOR signaling in human fetal neural stem cells, inhibiting neurogenesis and upregulating autophagy (Liang et al., 2016). In another study, by individually expressing each ZIKV protein in the mouse cortex *in vivo*, Yoon *et al*. discovered that NS2A inhibits neurogenesis and proliferation of neural stem cells by disrupting adherens junctions (Yoon et al., 2017). Protein interaction studies in human induced pluripotent stem cells found that Capsid interacts with Dicer, and further functional studies demonstrated that ZIKV inhibits Dicer and host micro RNA biogenesis which in turn disrupt neurogenesis (Zeng et al., 2020), By taking a systems level approach in humans to identify potential targets of ZIKV, we previous found that ZIKV protein NS4A physically interacts with ANKLE2, a protein that is encoded by a gene that is linked to a rare Mendelian form of congenital microcephaly (Khan et al., 2017; Yamamoto et al., 2014), in human cells (Shah et al., 2018). Using *Drosophila*, we further showed that expression of NS4A causes reduced brain volume *in vivo* which can be rescued by co-expression of reference human ANKLE2 (Link et al., 2019; Shah et al., 2018). These results show that NS4A binds to and inhibits ANKLE2, providing one compelling mechanism as to how ZIKV infection causes microcephaly. Particularly, this last study provides an example of how *Drosophila* can be utilized as a model system to elucidate mechanisms of infectious diseases (Harnish et al., 2021).

**Figure 1:**
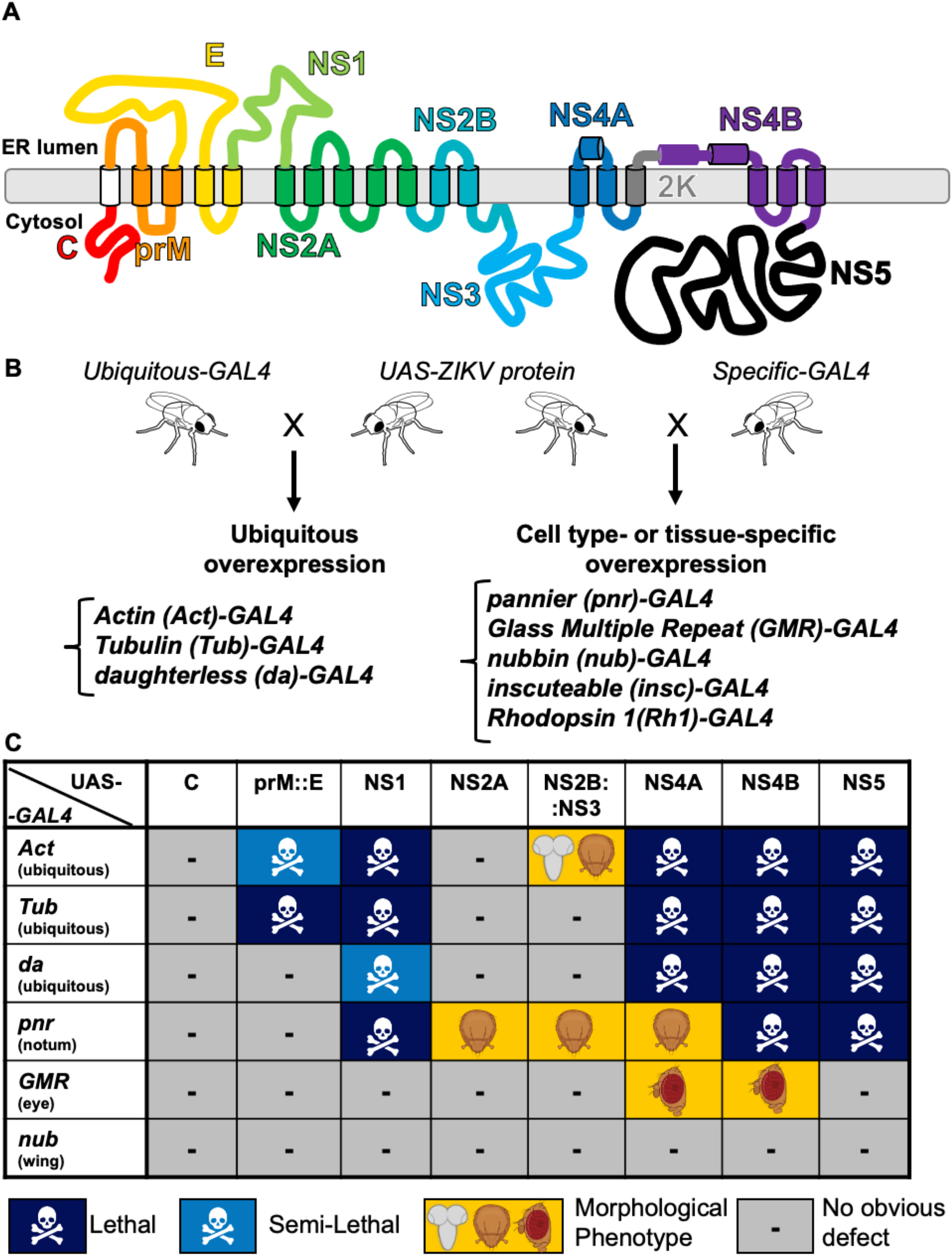
ZIKV proteins cause scorable phenotypes upon overexpression in *Drosophila*. A) Graphical diagram of the ZIKV polyprotein, which contains 10 proteins; 3 structural and 7 non-structural, at the ER membrane. B) Diagram of crossing scheme in which we cross a fly containing a ZIKV protein under the control of a UAS element to another fly containing a GAL4 driver. The resulting fly expresses the ZIKV protein and is scored for phenotypes. C) Table showing the phenotypes of the F1 generation resulting from a cross reared at 25 °C. Dark blue represents lethality. Light blue represents semi-lethality, where less than 75% of expected Mendelian ratios are observed. Yellow indicates a morphological defect; specific tissues affected are noted by the illustration (brain, eye, thorax bristle). Lethal stage for some crosses is indicated in **Supplementary Table 2**.

In this study, to better understand the mechanisms of ZIKV pathogenesis, we generated a comprehensive toolkit of transgenic flies that allow expression of ZIKV proteins using the GAL4/UAS binary expression system (Brand and Perrimon, 1993). In addition to generating transgenic lines that allow the expression of ZIKV proteins from the Puerto Rican strain associated with the recent epidemic of CZS (Lanciotti et al., 2016), we also generated several transgenic lines that allow expression of proteins from a less pathogenic viral strain isolated in Cambodia prior to the Zika virus epidemic (Liu et al., 2017; Xia et al., 2018; Yuan et al., 2017) to assess if the *Drosophila* resource can be used to study how virus evolution impacts pathogenesis. Using these tools and performing a series of phenotypic assays related to the nervous system, we show that *Drosophila* can deepen our understanding of ZIKV induced developmental and post-developmental neuronal symptoms in human.

## RESULTS

### Expression of most ZIKV proteins or protein complexes cause lethality or morphological phenotypes when ectopically expressed in Drosophila

To determine how ZIKV proteins might hijack host pathways to cause disease, we sought to express each protein in a wide variety of tissues throughout development and in the adult fly using the GAL4/UAS system (**Figure 1**). To generate transgenic flies, we used sequence information from the PRVABC-59 strain isolated in Puerto Rico (Lanciotti et al., 2016) and synthesized and cloned individual protein-coding sequences (following typical ZIKV polyprotein processing) into a UAS transgenic vector (*pGW-HA.attB*) (Bischof et al., 2013) using Gibson assembly (Gibson et al., 2009). Each protein of interest was tagged with a C’-3xHA tag for visualization and expression analysis, and careful consideration was taken to include proper signal sequences for transmembrane proteins (Shah et al., 2018). Transgenic strains were generated using the phiC31 transgenesis technique, allowing integration of all constructs into the identical genomic locus of the fly at the established *attP* docking site (VK37) at 22A3 on the second chromosome (Venken *et al*., 2006). We generated transgenic lines that allow expression of six proteins encoded in the ZIKV genome as single proteins: Capsid, NS1, NS2A, NS4A, NS4B, and NS5. We also generated a construct that combines prM with E (prM::E) and NS2B with NS3 (NS2B::NS3). The former transgene was generated because prM is known to act as a co-chaperone for proper folding of E, and the two proteins form heterodimers that are critical for flaviviruses to form infectious particles (Yu et al., 2008). The latter transgene was generated because NS2B and NS3 forms a heterodimer that functions as a key viral protease that processes the viral precursor polyprotein (Falgout et al., 1991). In addition, since there are proteolytic cleavage sites on both sides of the short peptide 2K located between NS4A and NS4B (Sun et al., 2017), we generated NS4A and NS4B with and without 2K (NS4A, NS4A::2K, 2K::NS4B and NS4B). We also generated NS1 and prM::E that alters specific amino acids in these proteins that have been implicated in infectability or pathogenicity. Specifically, we generated NS1^cam^ and prM::E^cam^, which carry single amino acid changes that are found in a less pathogenic strain of ZIKV, FSS13025, isolated in Cambodia (Liu et al., 2017). More specifically, NS1^cam^ carries an alanine (A) in position 188 of NS1 compared to the valine (V) in the Puerto Rican strain. We generated this transgene because the p.A188V mutation in NS1 has been found to increase the infectivity of ZIKV in mosquitos (Liu et al., 2017) and inhibit interferon-β production in human cells (Xia et al., 2018). prM::E^cam^ carries a serine (S) in position 139 of the E protein compared to the asparagine (N) the Puerto Rican strain. This construct was generated because a p.S139N mutation increased infectability of the virus in human and mouse neural progenitor cells and caused a more severe microcephaly phenotype in mouse models (Yuan et al., 2017). In total, we generated 12 transgenic lines (**Supplemental Table 1**).

First, we crossed eight transgenic lines (*UAS-C*, *UAS-prM::E*, *UAS-NS1*, *UAS-NS2A*, *UAS-NS2B::NS3*, *UAS-NS4A*, *UAS-NS4B*, *UAS-NS5*) from the Puerto Rican strain to a variety of GAL4 lines to express each viral protein or fusion proteins in a variety of tissues at different developmental time points to cast a wide net to assess host pathways that ZIKV might hijack during infection and disease progression. We tested three ubiquitous drivers with variable strengths [*tubulin(tub)-GAL4*, *Actin(Act)-GAL4*, and *daughterless(da)-GAL4*] and three tissue specific drivers [*nubbin(nub)-GAL4* for the wing, *pannier(pnr)-GAL4* for the dorsal thorax, and *GMR(Glass Multiple Reporter)-GAL4* for the eye] to assess the functional consequences of overexpressing the ZIKV proteins on viability and gross morphology. We found that seven out of eight lines (prM::E, NS1, NS2A, NS2B::NS3, NS4A, NS4B and NS5) caused scorable phenotypes with at least one of these drivers (**Figure 1C**, **Figure 2, Supplemental Table 1)**. Ubiquitous expression revealed that some viral proteins (prM::E, NS1, NS4A, NS4B and NS5) caused lethality, suggesting that these proteins likely impact essential cellular pathways. The lethal stage induced by some ZIKV proteins are presented in **Supplemental Table 2**. Overexpression of N2A caused increase in mechanosensory bristles, whereas NS2B::NS3 caused reduction of mechanosensory bristles on the dorsal thorax of the fly, indicating that these proteins affect pathways that are required for peripheral nervous system development (**Figure 1C, 2B-C**) (Schweisguth, 2015; Schweisguth et al., 1996). Overexpression of NS4A using *pnr-GAL4* caused a dorsal thorax closure defect (**Figure 1C, 2D**), which is a phenotype often seen when cell migration or cell communication through JNK and TGF-β/BMP signaling is defective (Agnès et al., 1999; Martin-Blanco et al., 2000). Overexpression of NS4A or NS4B using *GMR-GAL4* caused an rough eye morphology phenotype (**Figure 1C, 3E, 3G**), suggesting a developmental defect of the compound eye, which can be caused by disruption of many pathways and biological processes (Kumar, 2001; Thomas and Wassarman, 1999). In contrast, although the development of the wing depends on many signaling pathways and cellular processes (Bier, 2005), none of the lines examined, including the ZIKV proteins that cause lethality when overexpressed ubiquitously, caused an obvious wing morphological defect when expressed using *nub-GAL4*. Therefore, ZIKV proteins seem to affect specific proteins, pathways or cellular processes to induce a phenotype upon overexpression in flies in a context specific manner, rather than through inducing a general cellular toxicity.

**Figure 2:**
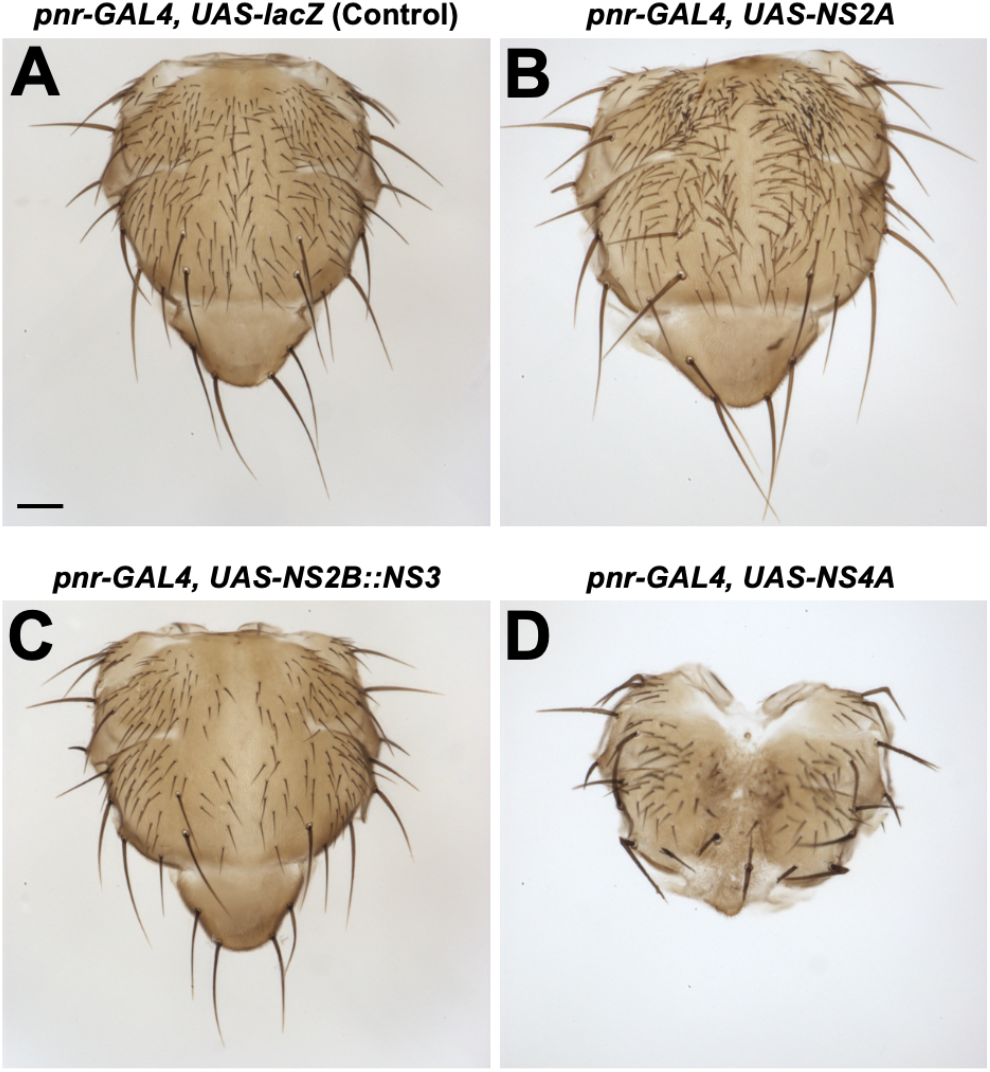
Expression of ZIKV proteins in the notum causes bristle and split thorax phenotypes. A) Control notum showing no morphological defects. B) Expression of NS2A in the dorsocentral notum causes a supernumerary bristle phenotype. C) Notum from animals with NS2B::NS3 expression (*pnr-GAL4, UAS- NS2B::NS3*) shows bristle loss. D) Rare escaper notum from animal with NS4A expression demonstrates a split thorax phenotype and bristle defects. All crosses were carried out at 29°C. Scale bar=0.1mm

### The site of 2K peptide cleavage can affect the function of NS4A and NS4B

Between the NS4A and NS4B proteins, there is a linker sequence that is referred to as the 2K peptide (**Figure 1A**) (Lin et al., 1993). The site between NS4A and 2K is cleaved by the viral NS2B::NS3 protease and the site between 2K and NS4B is cleaved by one or more host proteases, respectively (Sun et al., 2017). Because the two cleavage events occur sequentially (Lin et al., 1993) and 2K have been shown to alter NS4A function (Miller et al., 2007; Roosendaal et al., 2006), we tested whether expressing a fusion protein of NS4A and 2K (NS4A::2K) or 2K and NS4B (2K::NS4B) have different effect than expressing these two proteins without this peptide (**Figure 3A-B**).

**Figure 3:**
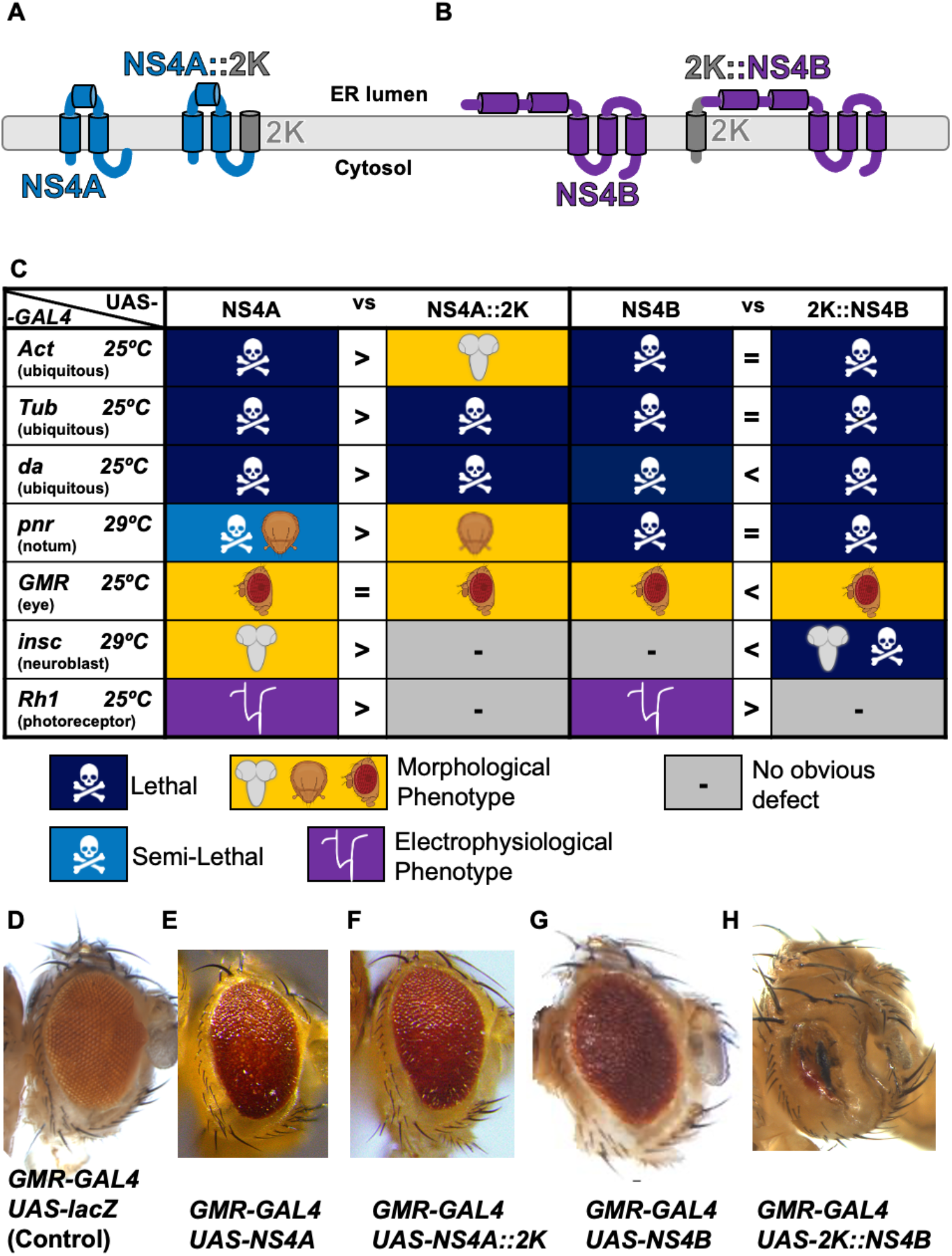
2K peptide alters NS4A and NS4B phenotypes. Graphical diagram of the ZIKV A) NS4A and NS4A::2K peptides or B) 2K::NS4B and NS4B peptides at the ER membrane. Both NS4A and NS4B can be found with the 2K peptide linker region. C) Table showing the phenotypes of the F1 generation resulting from a cross reared with indicated drivers at 25 °C or 29 °C. Dark blue represents lethality. Light blue represents semi- lethality, where less than 75% of expected Mendelian ratios are observed. Yellow indicates a morphological defect; specific tissues affected are noted by the illustration (brain, eye, thorax bristle). Lethal stage for some crosses indicated in **Supplementary Table 2**. (D-H) Eye phenotypes as a result of *GMR-GAL4* expression of control (D) lacZ, (E) NS4A, (F) NS4A::2K, (G) NS4B, and (H) 2K::NS4B. Note that in general, the 2K peptide decreases the effect of NS4A but can enhance the phenotypes caused by NS4B.

Ectopic expression of ZIKV proteins NS4A without the 2K peptide caused multiple scorable phenotypes with diverse GAL4 lines including lethality with *Act-GAL4*, *Tub-GAL4*, and *da-GAL4* (**Figures 1C and 3C**). Expression with *GMR-GAL4* produced animals with rough eye phenotypes (**Figures 1C, 3C and 3E**) and *pnr- GAL4* expression led to dorsal thorax closure (**Figure 1C, 2D and 3C**). When we expressed NS4A::2K with the same GAL4 drivers, we observed mostly weaker phenotypes compared to expression of NS4A alone (**Figure 3C and 3F**). Expression of NS4A caused lethality in most animals with few escapers using *pnr-GAL4* at 29°C, but lethality was not observed when NS4A::2K was expressed using this driver and temperature. However, ubiquitous expression of NS4A::2K with *da-GAL4* caused lethality during pupal stages, which is in contrast to NS4A causing early larval lethality using the same driver. Similarly, viable flies that express NS4A::2K using *Act-GAL4* were observed, which were not seen when NS4A was expressed using the same driver. In summary, the 2K peptide can sometimes suppress the function of NS4A depending on the temperature, context, and GAL4 drivers being used.

Similar to NS4A, ectopic expression of NS4B without the 2K peptide using multiple drivers affected viability and development. Expression of NS4B with all three ubiquitous drivers (*Act-GAL4*, *Tub-GAL4*, *da-GAL4*) caused lethality whereas expression in the developing eye using *GMR- GAL4* induced rough eyes (**Figure 1C, 3C and 3G**). We also observed that addition of 2K affected the severity of phenotypes with some drivers (**Figure 3C and 3H**). In three cases, the addition of 2K made the phenotype more severe. For example, expression of 2K::NS4B caused embryonic lethality whereas NS4B caused late larval lethality using *da-GAL4*. Similarly, expression of 2K::NS4B caused a much more severe eye developmental defect compared to NS4B using *GMR-GAL4* (**Figure 3C and 3G-H**). However, overexpression of 2K::NS4B did not change the lethality observed when NS4B was expressed using *Act-GAL4* or *Tub-GAL4*, and made the phenotype caused by another driver weaker (*Rh1-GAL4*, see below). Therefore, while addition of 2K peptide to NS4B enhances its function in some contexts, it can also be irrelevant or decrease its function in other circumstances.

### Multiple ZIKV proteins reduce brain volume when expressed in Drosophila

Since ZIKV infection is associated with severe microcephaly, we set out to define the set of viral proteins that might target important processes during brain development. Previously, we showed that expression of ZIKV NS4A::2K using ubiquitous (*Act-GAL4*) or neural stem cell and their progeny-specific [*inscutable(insc)-GAL4)*] drivers reduces the volume of third instar larval brains (Link et al., 2019; Shah et al., 2018). To determine whether other transgenes also have similar biological functions, we expressed each protein or fusion protein with *Act- GAL4* or *insc-GAL4*. Ubiquitous expression with *Act-GAL4* at 25°C, NS1, NS4A, NS4B, 2K::NS4B, and NS5 caused lethality prior to late third instar larval stage so brains could not be assessed (**Figure 4**). Expression of NS4A::2K using *Act-GAL4* caused significant brain size reduction as compared to controls (overexpressing CD8::GFP, a neutral transmembrane protein) (**Figure 4G, 4I**, Using estimation statistics (Ho et al., 2019), which uses bootstrap resampling to calculate effect size and determine confidence intervals, the unpaired mean difference between CD8::GFP and NS4A::2K is -1.42 [95.0%CI = -1.91, -0.965]. The *P* value of the two-sided permutation t-test is 0.0.), while expression of NS2B::NS3 caused a more mild but significant brain volume defect (**Figure 4F and 4I**, the unpaired mean difference between CD8::GFP and NS2B::NS3 is -0.838 [95.0%CI = - 1.33, -0.366]. All statistical values are presented in **Supplemental Table 3**). The *P* value of the two-sided permutation t-test is 0.0052). Using *insc-GAL4* at 29°C, we found expression of NS2B::NS3, NS4A, NS4A::2K, and 2K::NS4B caused significant brain volume defects (**Figure 4A-E, 4H**). In summary, these data not only support previous observations that NS4A expression (with or without 2K) in *Drosophila* causes reduced brain volume by interacting with the ANKLE2 pathway (rescue is shown in **Supplemental Figure 1**) (Link et al., 2019; Shah et al., 2018) but also identify NS2B::NS3 and 2K::NS4B as novel negative regulators of brain size when expressed in *Drosophila*.

**Figure 4:**
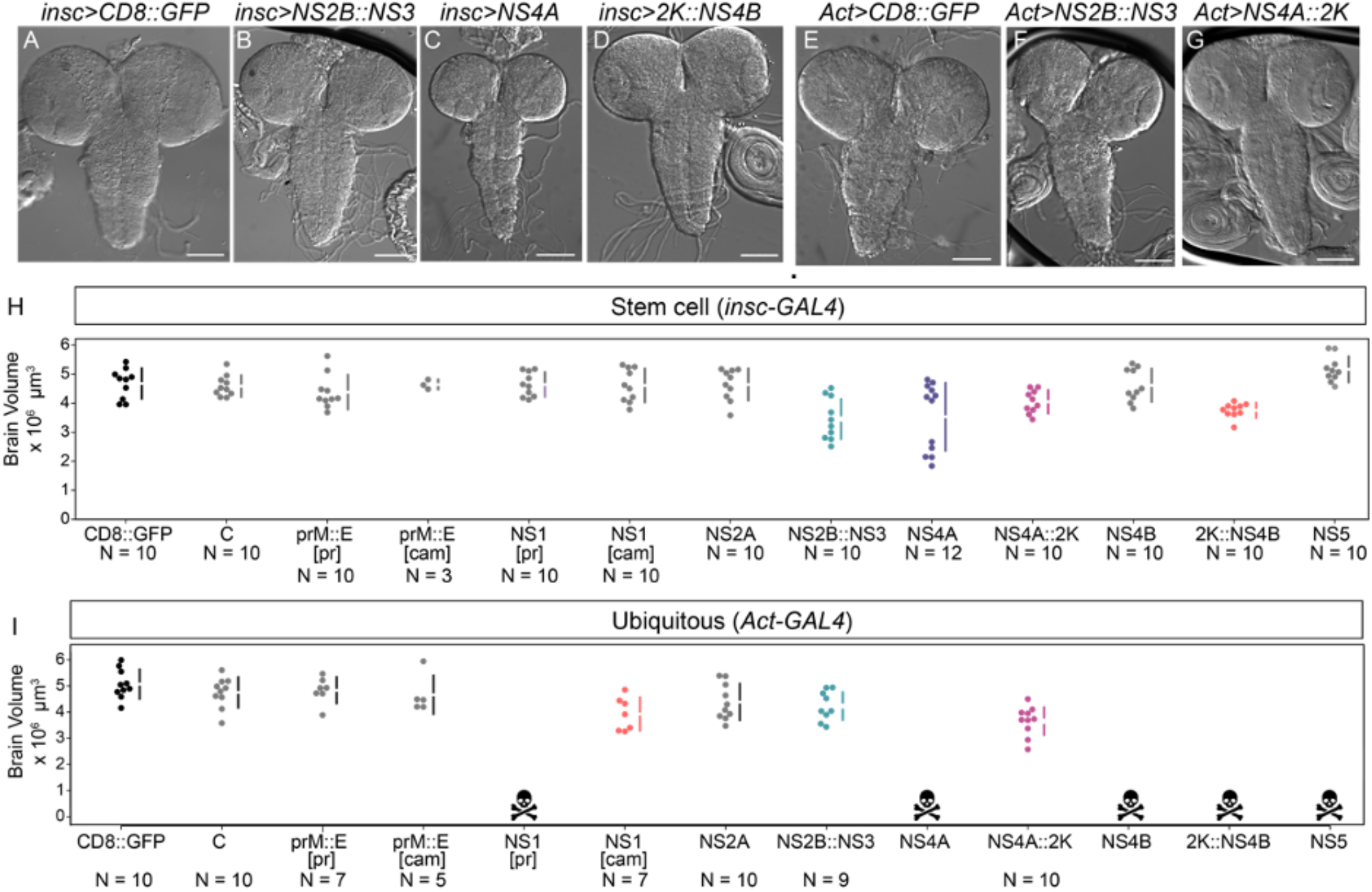
Multiple ZIKV proteins cause microcephaly upon overexpression in *Drosophila*. Expression of ZIKV proteins using either (A-D, H) *insc*-GAL4 at 29°C or (E-G, I) *Act*-GAL4 at 25°C causes microcephaly phenotypes. Bright field images (A-G) of indicated lines with brain volume quantified in (H) and (I). In (H-I), the Estimation Statistics online resource (Ho et al., 2019) was used to plot brain lobe volume measurements. The raw data is plotted in H and I with each individual measurement as a dot and the gapped lines represent the mean (gap) and standard deviations(lines). Effect size (mean difference), confidence intervals, and permutation t-test P values are located in Supplemental Table 3. Populations with smaller brain volumes and P values less than 0.05 are colored, while populations with P values higher than 0.05 are in grey.

### Immunostaining of epitope tags attached to ZIKV proteins expressed in Drosophila neuroblasts reveal different patterns of subcellular localization

ZIKV proteins are synthesized on rough endoplasmic reticulum (ER) and replication and assembly of the virus takes place primarily in this organelle (Cortese et al., 2017). However, there are several studies that suggest that some viral proteins may function in other subcellular organelles. For example NS5 can function in the nucleus to modulate host cell division and gene expression (Conde et al., 2020; Kesari et al., 2020; Li et al., 2022; Petit et al., 2021; Zhao et al., 2021). We assessed the expression and subcellular localization pattern of these proteins using immunofluorescence staining of the C’-3xHA tag and confocal microscopy. We dissected late third instar larval brains expressing a ZIKV transgene using *insc-GAL4* and immunostained them with Deadpan, a nuclear marker of neuroblasts (neural stem cells), and the HA tag, which labels each viral protein expressed. Many viral proteins were localized in a reticular pattern that was suggestive of ER localization (**Figure 5**), consistent with previous observations in mammalian cells (Hou et al., 2017). Some proteins exhibited distinct patterns, including NS4A and 2K::NS4B that showed a punctate pattern (**Figure 5J-K**) consistent with previous studies in mammalian cells (Miller et al., 2007; Roosendaal et al., 2006; Shah et al., 2018). NS5 showed nuclear localization (**Figure 5M**), also consistent with previous studies in mammalian cells (Petit et al., 2021). It is interesting to note that the presence or absence of 2K peptide on NS4A and NS4B, which can alter the phenotypic strength, causes a change in the subcellular localization of these proteins (**Figure 5I-L**) (**Figure 3**).

**Figure 5:**
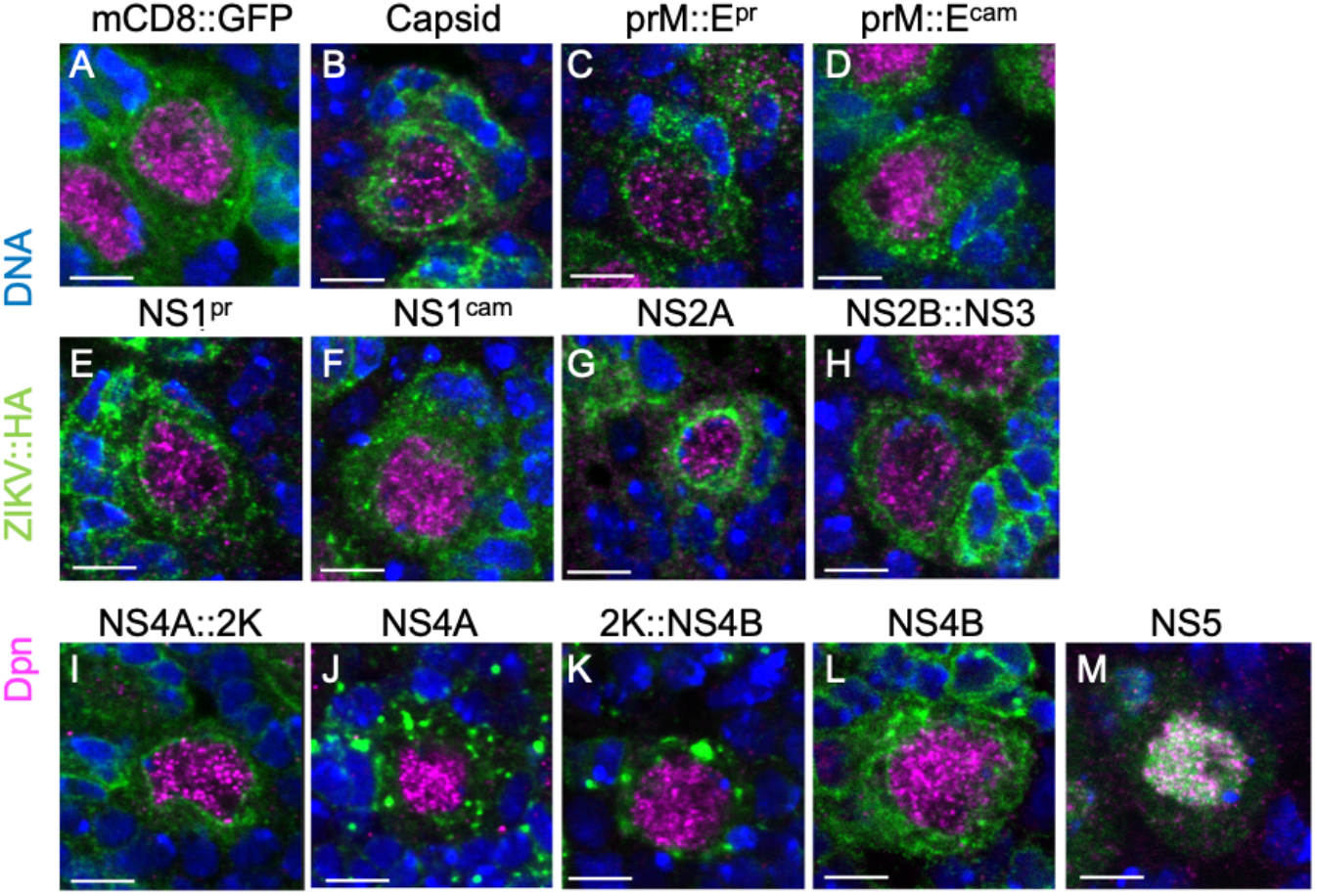
Subcellular localization of ZIKV proteins differ when expressed in neuronal stem cells. (A-M) Neural stem cells from third instar larvae with *insc*-GAL4 ZIKV protein expression stained with HA (green) to mark ZIKV proteins (tag is on the C-terminus), Dpn (purple) to indicate neuroblasts, and DAPI to highlight DNA. Each panel represents a single stem cell in interphase. Note that the 2K peptide alters protein localization of NS4A and NS4B while NS5 is localized in the nucleus.

### Expression of some ZIKV proteins in post-differentiated photoreceptor cells causes electrophysiological defects

Adult humans infected with ZIKV can present with neurological symptoms (Mehta et al., 2018), while adult mice exposed to this virus can exhibit long-term neuropathological defects (Nem de Oliveira Souza et al., 2018). Thus, it is possible that ZIKV may infect and impact the function of the post-developmental nervous system. To test whether ZIKV proteins affect post-differentiated neurons, we expressed each protein in the fly photoreceptor cells using *Rhodopsin 1(Rh1)*-GAL4 at 25°C. Photoreceptors in insects are light-sensory neurons that project axons into the lamina or medulla layers of the adult brain. *Rh1* [also known as *ninaE (neither inactivation nor afterpotential E)*] encodes a Rhodopsin that is expressed in six out of eight photoreceptors (R1 to R6) that comprise an ommatidia, the basic unit of a compound eye. The function of photoreceptor neurons can be assessed using electroretinogram (ERG) analyses, which are a cumulative measure of neuronal function in response to light from the *Drosophila* eye and brain (Ugur et al., 2016). When light is flashed on, an ERG trace that consists of an on-transient spike followed by depolarization is recorded. When light is turned off, an off-transient spike is observed, followed by repolarization of the potential to baseline (**Figure 6**). Depolarization/repolarization represents the phototransduction response of photoreceptors to light stimulus, while the on/off transient spikes correspond to signal transmission to between pre- and post-synaptic neurons in the brain (Dolph et al., 2011). Adult animals expressing each viral protein were aged for 30 days post-eclosion in a 12 hour light/dark cycle at 25°C. For most ZIKV proteins, expression in photoreceptor cells did not affect ERG patterns. However, prM::E expression caused a strong decrease in ERG depolarization amplitude, indicating photoreceptor function defects (**Figure 6B, D**). In addition, expression of NS4A, but not NS4A::2K, caused loss of ERG on/off transient spikes in aged animals (**Figures 3C, 6C, E, F, J-L**). To determine whether this phenotype is age-dependent, we tested the ERG on flies expressing NS4A using *Rh1-GAL4* that were only aged for 5 days post-eclosion. Interestingly, we did not observe a difference in the on/off transient spikes or depolarization in these young animals (**Figure 6G-I**). Therefore, the ERG defect caused by expression of NS4A is likely to be degenerative rather than developmental or physiological in nature. Finally, Expression of NS4B, but not 2K::NS4B mildly decreased on transient amplitude. Importantly, expression of other ZIKV proteins did not show these phenotypes, indicating that prM::E, NS4A, and NS4B specifically inhibit the function and activity of mature neurons.

**Figure 6:**
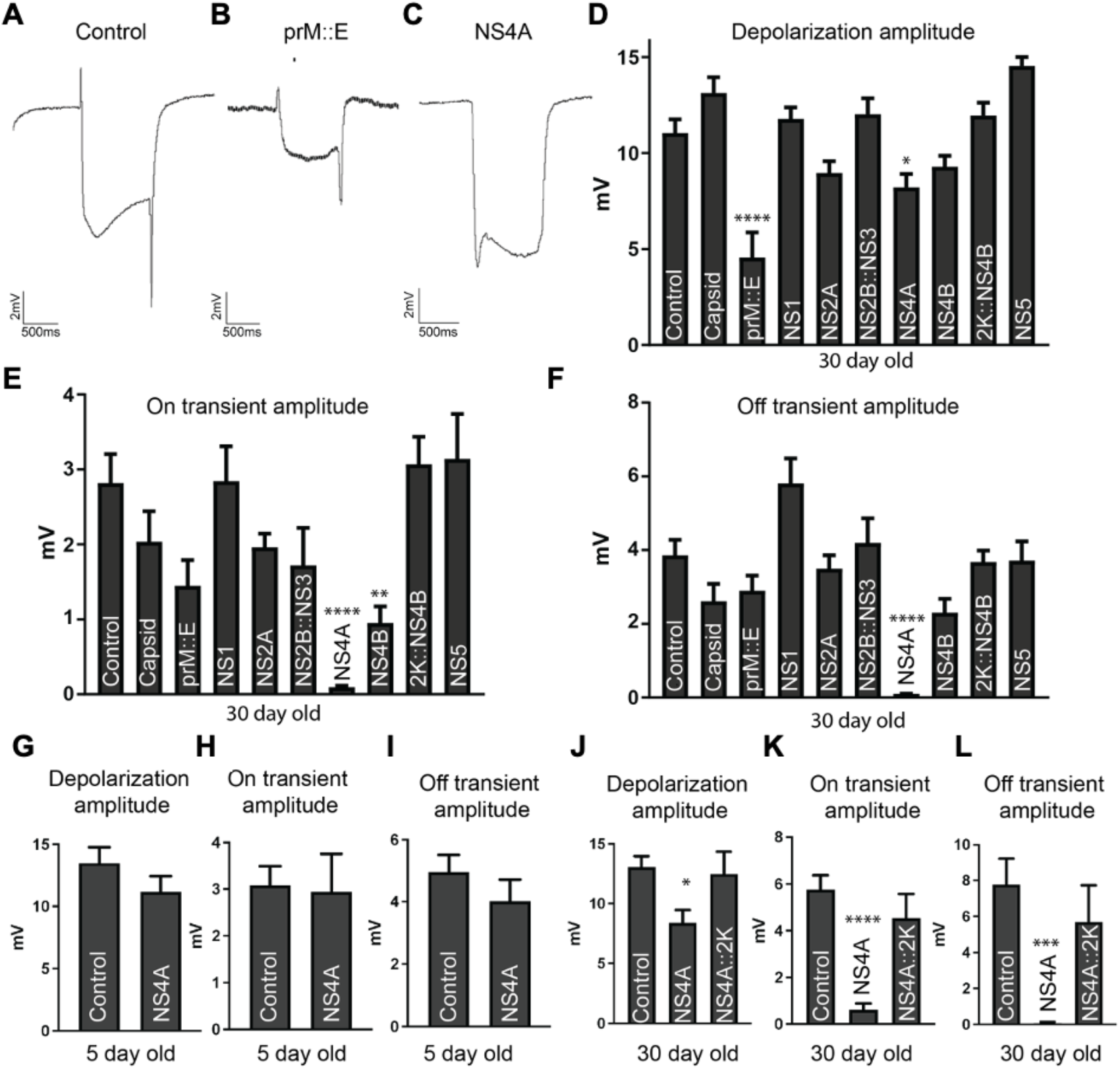
Expression of some ZIKV proteins causes electrophysiological defects in the fly visual system. (A-C) Representative electroretinograms from (A) control (luciferase), (B) prM::E, and (C) NS4A expressing animals 30 days after eclosion in 12 hr light/dark cycle . Note reduced depolarization amplitude in animals with neuronal expression of prM::E, quantified in (D), and loss of on- and off-transients with NS4A expression, quantified in (E) and (F). (G) ERG depolarization amplitude, (H) on-, and (I) off- transient quantification of NS4A expressing animals at 5 days after eclosion. No defect is documented, indicating NS4A causes degenerative ERG defects over time. (J-L) Comparisons between NS4A alone or NS4A::2K show that only NS4A alone causes neuronal phenotypes at 30 days after eclosion.

### ZIKV proteins encoded by different strains show functional differences when tested in Drosophila

Finally, we explored if the *Drosophila* system could be utilized to test the functional consequences of mutations that were acquired in protein coding genes during ZIKV evolution. The more recent Puerto Rican ZIKV strain, which is more closely related to strains responsible for recent outbreaks in the Americas, causes high rates of fetal microcephaly, similar to outbreaks in French Polynesia (Cauchemez et al., 2016). However, before these outbreaks, ZIKV infection was not associated with microcephaly. Since the Cambodian strain was the last virus of the Asian lineage not associated with severe microcephaly (Duong et al., 2017), it is useful for comparative investigations. We focused on two mutations, one in NS1 and the other in E, as they had both been previously suggested to contribute to disease severity using other experimental systems (Wang et al., 2017; Xia et al., 2018; Yuan et al., 2017).

The NS1 protein from the Puerto Rican strain (NS1^pr^, which is the protein tested in assays mentioned above) carries a p.A188V mutation compared to NS1 protein from the more ancient Cambodian strain (NS1^cam^) (Delatorre et al., 2017). This mutation has been found to enhance the evasion of ZIKV from the host immune system based on studies performed in mice and human cells (Xia et al., 2018). When expressed using *Act-GAL4*, *Tub-GAL4* or *pnr-GAL4*, NS1^pr^ caused lethality (**Figure 1C, 7A**). However, expression of NS1^cam^ was only semi-lethal using the same drivers tested under the same condition (**Figure 7A**). This suggests that NS1^pr^ has stronger function compared to NS1^cam^, consistent with the notion that the Puerto Rican isolate is more pathogenic than the Cambodian isolate.

**Figure 7:**
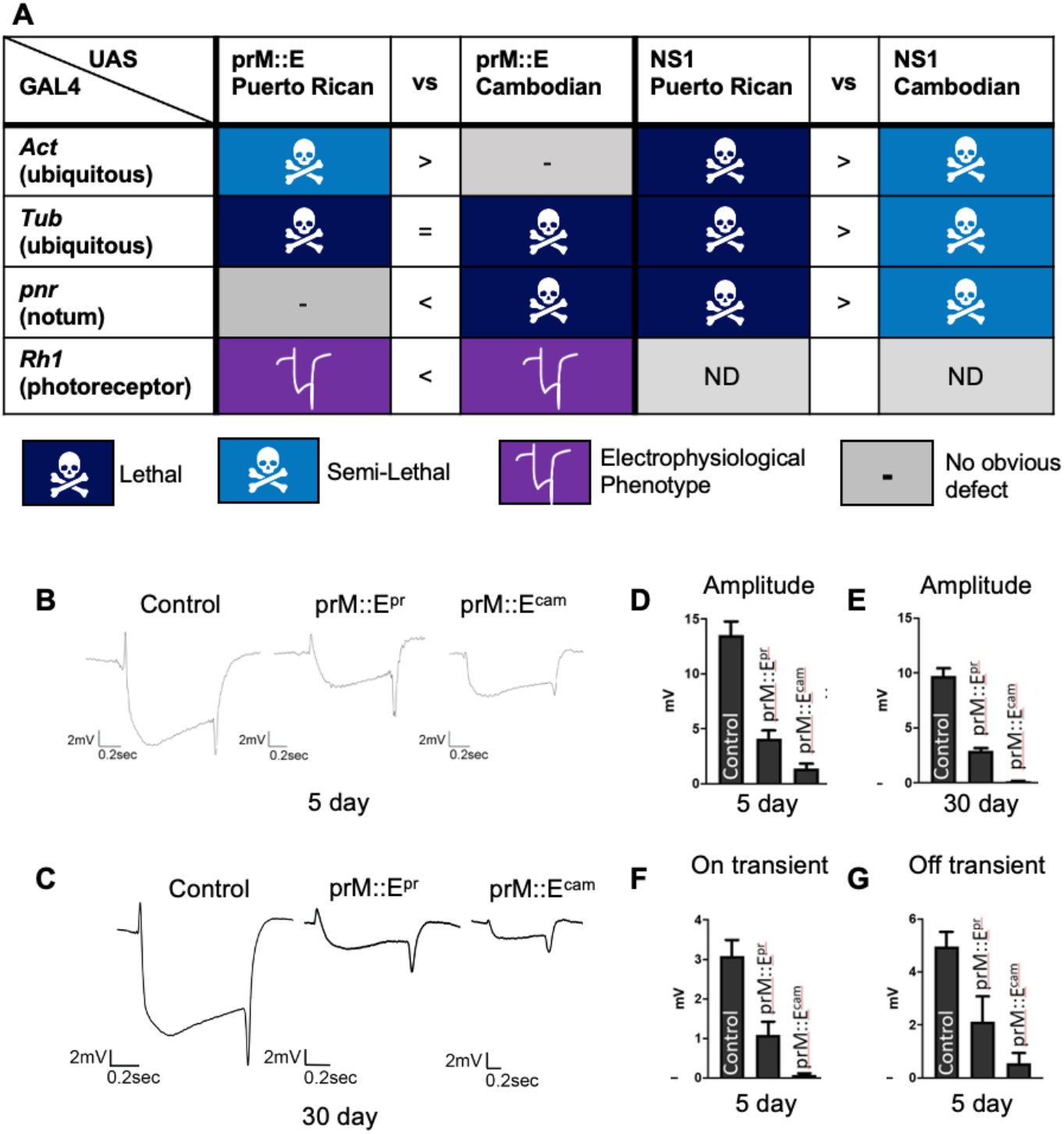
ZIKV protein expression phenotypes from prM::E and NS1 are different based on viral strain. A) Phenotypes when prM::E and NS1 from either the Puerto Rican or Cambodian strains of ZIKV are expressed by various GAL4 drivers. “Vs” column denotes if one variant is more severe with > or <. = denotes equal severity. (B) Representative ERG traces of control, prM::E^pr^, and prM::E^cam^ at 5 days or (C) 30 days after eclosion. (D) Depolarization amplitude of 5 day or (E) 30 day old animals expressing luciferase (control), prM::E^pr^, or prM:E^cam^ shows developmental ERG defects that are stable with age. prM::E^cam^ is more severe than prM::E^pr^. (F) On transient and (G) off transient amplitudes are also reduced with expression of prM::E^pr^ or prM:E^cam^ shown at 5 days after eclosion. ND=not determined.

The E protein from the Puerto Rican strain (prM::E^pr^, which is the protein tested in assays mentioned above) carries a p.S139N mutation compared to NS1 protein from the Cambodian strain (prM::E^cam^). This mutation has been associated with increased infectivity and severity of microcephaly when tested in mice and human cells (Yuan et al., 2017). prM::E^pr^ caused semi-lethality when expressed with *Tub-GAL4* (**Figure 1C and 7**) and exhibited an ERG defect when expressed with *Rh1-GAL4* (**Figure 6**). While we did not observe a difference between prM::E^pr^ and prM::E^cam^ when these proteins were expressed using *Tub-GAL4*, we identified functional differences when *pnr-GAL4* and *Rh1-GAL4* were used. In contrast to prM::E^pr^ that does not show any defect when driven with *pnr-GAL4*, prM::E^cam^ expression using this driver caused lethality (**Figure 7A**). Similarly, expression of prM::E^cam^ caused a stronger ERG phenotype compared to prM::E^pr^ using Rh1-GAL4 (**Figure 6B-E**). These data indicate that the prM::E^cam^ is more deleterious compared to prM::E^pr^, which was opposite from what we expected. Regardless, these data show that the *Drosophila* assay system can be used to identify functional differences of mutant viral proteins.

## DISCUSSSION

We developed new transgenic *Drosophila* strains that allow researchers to investigate how viral proteins affect host pathways to cause disease. We expressed all ten ZIKV proteins (some as fused proteins) in eight contexts using three ubiquitous and five tissue specific GAL4 drivers. With the exception of Capsid, we were able to identify scorable phenotypes for all viral proteins, either when expressed alone or when co-expressed with their known functional partners. These phenotypes are not uniform; each viral protein shows different phenotypes in diverse contexts, suggesting that each protein interferes with specific developmental and cellular processes in distinct tissues.

ZIKV infection has been linked to severe microcephaly (de Araujo et al., 2018), a devastating phenotype leaving the patient dependent on others throughout their life. Previous reports using a comparative proteomics approach and a *Drosophila* model of microcephaly demonstrated that NS4A expression affects brain development by inhibiting the ANKLE2 pathway (Link et al., 2019; Shah et al., 2018). The results presented here verify this observation and emphasize the reliability of genetic model systems such as the fruit fly. By testing additional ZIKV proteins in the same microcephaly model, we demonstrated that NS2B::NS3 and NS4B also likely contribute to this phenotype. Host pathways disrupted by these proteins may unravel additional mechanisms by which ZIKV causes microcephaly. Furthermore, established ZIKV-human proteomic interaction data set (Shah et al., 2018) will be valuable to illuminate important host targets of these proteins to provide mechanistic insights into ZIVK associated diseases for therapeutic development.

During ZIKV infection, each viral protein is expressed at high levels which may interfere with multiple host pathways simultaneously. Combinatorial expression of multiple ZIKV proteins may also provide insight as to how hijacking of multiple developmental pathways coalesces to result in the diversity of established patient outcomes. Given that NS2B::NS3, NS4A, and NS4B all give rise to developmental phenotypes in the brain, co-expression of these proteins may result with more severe phenotypes which warrants future exploration. We also noted that the function of NS4A and NS4B can be significantly altered by the presence or absence of the small peptide 2K. Cleavage at the 2K site in different flaviviruses has been linked to membrane rearrangements induced during flavivirus replication (Miller et al., 2007; Roosendaal et al., 2006) or changes in flavivirus protein function (Plaszczyca et al., 2019). Interestingly, the subcellular localization of NS4A::2K and 2K::NS4B differed from NS4A and NS4B alone, respectively. It is possible that the 2K cleavage is an important regulatory step during viral replication and phenotype development, possibly modulating the function of these proteins.

Comparative analysis of proteins from the Puerto Rican or Cambodian ZIKV strains gave interesting insights into ways in which *Drosophila* can be used to study viral evolution. Modern ZIKV infection is associated with fetal microcephaly, however earlier strains of the virus presented have not been associated with brain developmental defects (Chu et al., 2017; Duong et al., 2017; Kuadkitkan et al., 2020). Our experiments in which we expressed both the Puerto Rican and Cambodian strains suggest that NS1^pr^ may have more severe effects on the host. However, prM::E^pr^ expression resulted in more milder phenotypes than prM::E^cam^ expression, as per experiments performed using two GAL4 drivers (*Rh1-GAL4* and *pnr-GAL4*) with two distinct phenotypes (ERG and lethality). This was unexpected as the p.S139N variant present in prM::E^pr^ has been reported to worsen the infection outcome in mice (Yuan et al., 2017). These discrepancies in data may represent a difference in viral effects and mechanisms of action in different tissues. Nevertheless, considering that we observed a phenotypic difference in both cases examined, *Drosophila* could be used as a primary screening system to identify mutations that have a functional consequence that can be further tested in cellular models or rodent systems.

Finally, our results show that *Drosophila* can be used as a system to explore the post-developmental effects of viral proteins on neuronal function. The *Drosophila* eye is a well-established model to investigate neural function and degenerative phenotypes that are often associated with disease (Ugur et al., 2016). While expression of ZIKV NS4A caused no obvious phenotypes in young flies, aged animals showed significant reduction in ERG transient amplitudes. Our data suggest that neurons expressing NS4A are functional early in life but may become less functional or degenerate over time. These results implicate that late-onset neuronal dysfunction might be a concerning phenotype following ZIKV infection. Indeed, studies in mouse and human iPSC (induced pluripotent stem cell)-based studies also found that exposure to ZIKV can result in neurodegeneration or make neurons more susceptible to insults (Bellmann et al., 2019; Noguchi et al., 2020). Hence, it will be imperative to follow how neuronal function changes after ZIKV infection in children and adults.

## ACKNOWLEDGMENTS

This study was supported in part through the Baylor College of Medicine Junior Faculty Seed Funding Award to S.Y. from the Naman Family Fund for Basic Research and Caroline Wiess Law Fund for Research in Molecular Medicine. B.H. was funded in-part by the Baylor College of Medicine Postbaccalaureate Research Education Program NIGMS grant R25GM069234. Confocal microscopy was supported in part by the NICHD grant U54HD083092 to the Intellectual and Developmental Disabilities Research Center (IDDRC) Neurovisualization Core at Baylor College of Medicine. N.L. and P.S.S. were funded in-part by NIAID grant R56AI170857 and R01AI170857.

## EXPERIMENTAL MODEL AND SUBJECT DETAILS

### Drosophila melanogaster strains and culture

The following fly lines were used: *PBac{y[+mDint2] w[+mC]=UAS-Capsid^pr^.HA*}*VK00037* (this study), *PBac{y[+mDint2] w[+mC]=UAS-prM::E^pr^.HA}VK00037* (this study), *PBac{y[+mDint2] w[+mC]=UAS- prM::E^cam^.HA}VK00037* (this study), *PBac{y[+mDint2] w[+mC]=UAS-NS1^pr^.HA}VK00037* (this study), *PBac{y[+mDint2] w[+mC]=UAS-NS1^cam^.HA}VK00037* (this study), *PBac{y[+mDint2] w[+mC]=UAS- NS2A^pr^.HA}VK00037* (this study), *PBac{y[+mDint2] w[+mC]=UAS-NS2B::NS3^pr^.HA}VK00037* (this study), *PBac{y[+mDint2] w[+mC]=UAS-NS4A^pr^.HA}VK00037* (this study), P*Bac{y[+mDint2] w[+mC]=UAS- NS4A::2K^pr^.HA}VK00037* (this study), *PBac{y[+mDint2] w[+mC]=UAS-2K::NS4B^pr^.HA}VK00037* (this study), *PBac{y[+mDint2] w[+mC]=UAS-NS4B^pr^.HA}VK00037* (this study), *PBac{y[+mDint2] w[+mC]=UAS- NS5^pr^.HA}VK00037* (this study), *P{UASt-CD8-GFP}* (Lee and Luo, 2001), *Act-GAL4* (*P{Act5C-GAL4}17bFO1*)(Ito et al., 1997), *inscuteable-GAL4* (*P{w[+mW.hs]=GawB}insc[Mz1407]*) (Luo et al., 1994), and *daughterless-GAL4* (*P{w[+mW.hs]=GAL4-da.G32}UH1*) (Wodarz et al., 1995). All flies were maintained at room temperature (∼22°C) and grown on standard cornmeal and molasses medium in plastic vials. Crosses were performed at temperatures indicated (25°C or 29°C). All studies contained male and female flies. Brain volume measurements were conducted in late 3rd instar larvae (judged by gut clearance and extruding spiracles), and all other assessments occurred in adults.

## METHODS

### Generation of transgenic constructs

To generate ZIKV expression constructs, ZIKV sequence was acquired from the strain PRVABC-59 (GenBank: KX377337.1), an Asian lineage isolated from Puerto Rico in 2015. Constructs were designed as described in (Shah et al., 2018). All sequences were codon optimized for expression in *Drosophila* using IDT’s Codon Optimization Tool (https://www.idtdna.com/CodonOpt). For initial fragments (Capsid, prM::E, NS1, NS2A, NS2B::NS3, NS4A, 2K::NS4B, NS5), double stranded, linear gene fragments were generated using IDT gBlocks. ZIKV fragments were synthesized with additional regions of homology that correspond to sequences in the destination vector (pGW-HA.attB) at the 5’ (5’-tcaaaaggtataggaacttcaaccggt<u>caac</u>-3’, Kozak sequence underlined) and 3’ (5’-gtacctcgaagttcctattctctacttagtata-3’) ends. AgeI and KpnI restriction enzymes were used to remove the gateway compatible cassette from pGW-HA.attB, removing attR1, CmR, ccdB, and attR2 regions. UAS expression constructs were generated by combining the linearized pGW-HA.attB backbone and each individual ZIKV fragment in Gibson assembly reactions (NEB Gibson Assembly Cloning Kit, E5510S). To generate NS4A::2K, NS4A and the 2K peptide (with 25bp overlapping regions included in primers) were PCR amplified, gel purified, and used in a 3 fragment Gibson assembly reaction. To generate the NS4B construct without the 2K peptide, NS4B was PCR amplified, gel purified, and assembled with pGW-HA using Gibson assembly. To generate prM::E and NS1 containing single amino acid changes seen in a Cambodian strain (FSS13025), variant products were synthesized and cloned into pGW-HA using Gibson assembly as above. All constructs were inserted into the VK37 landing site on the 2nd chromosome in *Drosophila melanogaster* by microinjection into a strain expressing ɸC31 integrase in the germline (*y[1] M{RFP[3xP3.PB] GFP[E.3xP3]=vas- int.Dm}ZH-2A w[*]; PBac{y[+]-attP-3B}VK00037*) (Bischof et al., 2007; Venken et al., 2006). Transgenic lines were selected by the *white^+^* marker in a *white^-^* background, and stable stocks were established through standard procedures (Harnish et al., 2019).

### Crossing Schemes and scoring of gross phenotypes

Wing, eye and thorax phenotypes were explored by driving the *UAS-ZIKV cDNA* lines with *nub-*GAL4*, GMR-Gal4 and pnr-Gal4*, respectively. Virgin females from the GAL4 stocks were collected from stock bottles kept at 25°C. Male flies were collected from the UAS stocks. Virgin females were isolated for 24 hours before being crossed, and 3-4 females to 3-5 males were placed in the same vial containing standard cornmeal and molasses medium. Crosses were initially maintained at 25°C. The crosses were transferred into new vials after 4-6 days and maintained at 25°C. After 4-6 days these parental flies were again transferred to new vials and maintained at 29°C. New flies were added if there were an insufficient number for the 29°C cross. Flies were collected 6-8 days after flies had begun to eclose. F1 progeny were scored for gross morphological phenotypes after being anesthetized with carbon dioxide. If phenotypes were observed, the flies were sacrificed via overnight freezing at -20°C and imaged the following day. The wings were dissected dry in a siliconized dish and imaged at 5x magnification. To assess lethality, the rate of flies carrying balancer chromosomes in the F1 generation was compared to the expected rate to determine whether flies were eclosing according to Mendelian ratios. Semi-lethality was noted when the observed number of protein-expressing flies was less than 75% of expected based on mendelian ratios.

### Immunostaining of larval brains

Late third instar (judged based on gut clearance and extruding spiracles) larval brains were dissected in PBS and fixed with 4% PFA/PBS/0.3% Triton for 20 minutes. For immunostaining, brains were blocked in PBS/0.3% Triton/1% BSA/5% normal goat serum and incubated in primary antibody in PBS/0.3% Triton/1% BSA overnight. Primary antibodies include rat anti-Deadpan (Abcam Cat# ab195172, 1:250 or 1:500), mouse anti-Prospero (Developmental Studies Hybridoma Bank, MR1A, 1:1000), and mouse anti-HA (Covance Cat# 901501, 1:1000) with donkey secondary antibodies from Jackson ImmunoResearch used at 1:500. Brains were mounted with double sided tape spacers and imaged using a confocal microscope (Leica Sp8) with 2 µm or 3 µm sections through the entire brain lobe.

### Brain volume measurement

Brains from third instar larvae were stained and mounted with tape spacers and imaged using a Leica Sp8 with 2 µm or 3 µm sections through the entire brain lobe. Resulting stacks were analyzed using the Surfaces function in Imaris (Bitplane) to quantify brain lobe volume as total microns cubed. One lobe from each brain was imaged and a total of 10 brains were analyzed per genotype or condition. Brain lobe volumes are displayed as box plots with hinges representing the 25th to 75th percentiles, a line represents the median, and whiskers represent min to max. Statistical significance was determined using one-way ANOVA with multiple comparisons post-test calculated using GraphPad Prism. Brain volumes are displayed as total brain volume (μm^3^).

### Western blot analysis

One to five brains from third instar larvae expressing *UAS-ZIKV cDNA* constructs with *Insc-GAL4* were were dissociated in 0.1% CHAPS buffer (50mM NaCl, 200mM HEPES, 1mM EDTA, Roche) supplemented with protease and phosphatase inhibitors for at least 30 min on ice, centrifuged for 10 min at 4°C, and supernatant was used for western analysis. Primary antibodies include mouse anti-HA (Covance Cat# 901501, 1:1000). Secondary antibodies include Rockland DyLight 600 or 800 (1:1000). Blots were imaged on a Bio-Rad ChemiDocMP.

### Electroretinogram recordings

*Rh1*-GAL4 was crossed to each ZIKV expression line, progeny were collected and aged at 25°C with a 12hr light/dark cycle (500 Lux) for 30 days. ERG recordings were performed as follows: flies were immobilized on glass slides with Elmer’s glue, a glass reference electrode filed with 100mM NaCl was inserted behind the eye, and a sharp recording electrode filled with 100mM NaCl was placed on the eye. Light flashes of 1 second followed by 1 sec recovery were delivered using a halogen lamp. 8-10 flies were tested for each genotype. Data was recorded and analyzed using AxoScope pClamp.

## SUPPLEMENTAL FIGURES

**Supplemental Figure 1:**
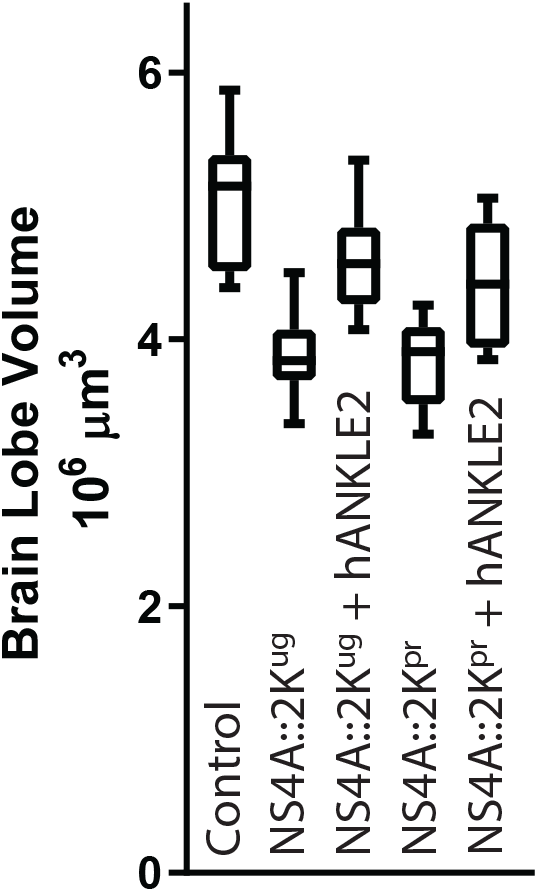
Human ANKLE2 rescues NS4A::2K induced defects from both Puerto Rican and Ugandan strains. Brain volume of third instar larvae with expression of NS4A::2K^ug^ (Uganda) or NS4A::2K^pr^ (Puerto Rico) with or without wild type human *ANKLE2*, which rescues NS4A::2K induced microcephaly.

**Supplemental Table 1:**
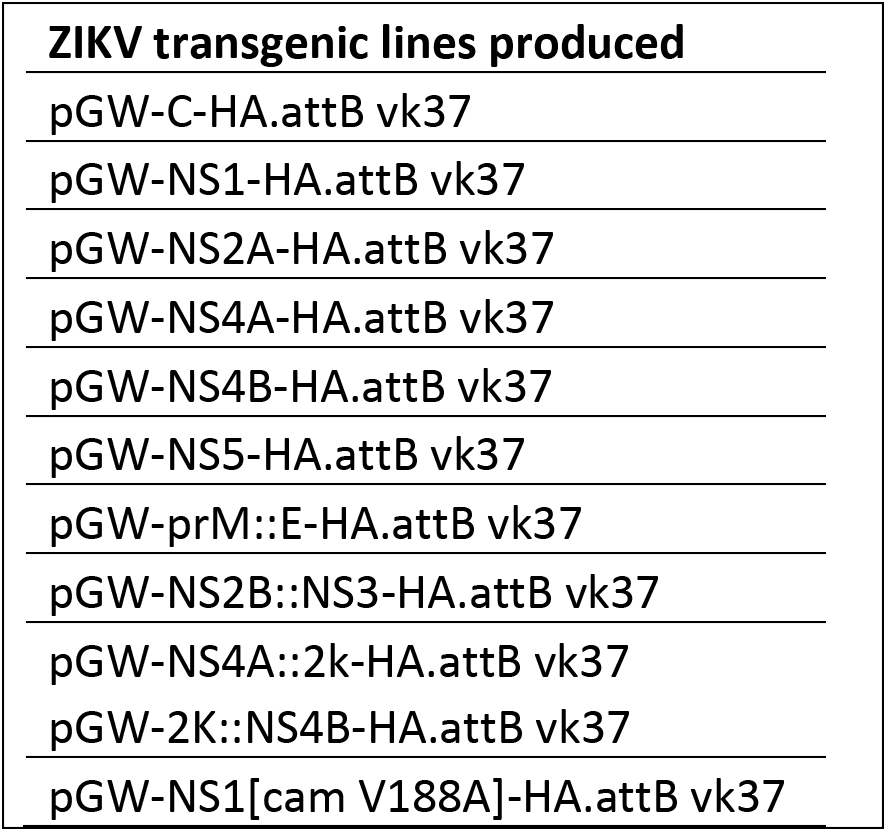
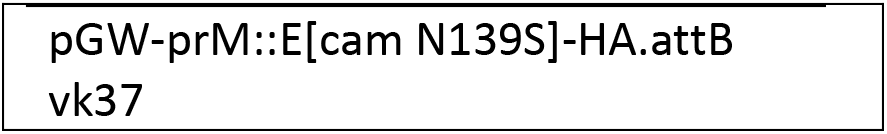
A list of all ZIKV transgenic lines produced. List includes backbone vector and attP site.

| ZIKV transgenic lines produced |
| --- |
| pGW-C-HA.attB vk37 |
| pGW-NS1-HA.attB vk37 |
| pGW-NS2A-HA.attB vk37 |
| pGW-NS4A-HA.attB vk37 |
| pGW-NS4B-HA.attB vk37 |
| pGW-NS5-HA.attB vk37 |
| pGW-prM::E-HA.attB vk37 |
| pGW-NS2B::NS3-HA.attB vk37 |
| pGW-NS4A::2k-HA.attB vk37 |
| pGW-2K::NS4B-HA.attB vk37 |
| pGW-NS1[cam V188A]-HA.attB vk37 |

**Supplemental Table 2:** Lethal stage of ZIKV crosses. GAL4 driver line and transgenic lines are noted. If animals were lethal prior to eclosion, the lethal stage is noted. ND = not determined.

| **GAL4** | **C** | **prM::E** | **NS1** | **NS2A** | **NS2B::NS3** | **NS4A** | **NS4A::2K** | **2K::NS4B** | **NS4B** | **NS5** |
| --- | --- | --- | --- | --- | --- | --- | --- | --- | --- | --- |
| *Act* | - | pupal | L1 | - | - | L2 | - | embryonic | embryonic | ND |
| *Tub* | - | ND | ND | - | - | before L3 | pupal | early L3 | before L3 | ND |
| *da* | - | - | escapers | - | - | L1 | pupal | embryonic | L3 | L2 |
| *pnr* | - | - | ND | - | - | escapers | - | embryonic | embryonic | ND |
| *GMR* | - | - | - | - | - | - | - | - | - | - |

**Supplemental Table 3: Estimation statistics for brain volume. Estimation statistics for brain volume analysis (related to** **Figure 4**). Estimation statistics, which focuses on effect size, were used to analyze brain volume data. 5000 bootstrap samples were taken; the confidence interval is bias-corrected and accelerated. The *P* value(s) reported are the likelihood(s) of observing the effect size(s), if the null hypothesis of zero difference is true. For each permutation *P* value, 5000 reshuffles of the control and test labels were performed.

